# Temperate versus virulent phage lifestyle impacts microdiversity in soil environments

**DOI:** 10.64898/2025.12.19.695463

**Authors:** Thomas E.P. de Bruijn, Hilje M. Doekes, Anne Kupczok

## Abstract

Bacteriophages are not only the most ubiquitous biological entity on earth, they also display remarkable genetic diversity across and within populations. While macrodiversity has been extensively studied, the drivers of microdiversity (intra-species genetic diversity) remain poorly understood, particularly in relation to phage lifestyle. The distinguishing ability of temperate phages to integrate themselves into the host genome has an unknown influence on the microdiversity present. This difference in microdiversity could impact the adaptability of phages to (a)biotic factors. To identify a possible association between microdiversity and lifestyle, we analysed 12 existing viromics datasets focusing on soil bacteriophages, including 43 171 viral genomes in total. We found that phages predicted to be temperate consistently exhibit higher microdiversity than their virulent counterparts in eight of 12 datasets. Of the remaining four datasets, one showed the opposite trend and three datasets did not show a significant trend at all. The detected pattern holds across multiple quality thresholds and lifestyle prediction methods. These findings suggest that lifestyle has a significant impact on genetic variation within phage populations, with potential implications for phage-host coevolution and environmental adaptability.

## 1. Introduction

Viruses are known to be the most ubiquitous biological entities on Earth, exerting immense influence over prokaryotic and eukaryotic life. Bacteriophages for example shunt and shuttle nutrients throughout ecosystems by lysing up to 40% of marine bacterial populations each day [1, 2, 3, 4]. The ubiquity of bacteriophages is reflected in—and perhaps due to—their morphological and genetic diversity [5, 6]. Advances in sequencing methods have enabled researchers to study this genetic diversity of bacteriophages at an unprecedented scale and precision [1].

Recent research into microbiome biodiversity has unearthed the vast diversity of bacteriophages, especially in soil environments [7, 2, 8, 9]. Historically, this biodiversity is described as macrodiversity or any of the many related terms (e.g. species richness, *α/β/γ*-diversity)[10]. The decreased costs and improved accuracy of DNA sequencing have also paved the way to study another type of biodiversity, the so-called microdiversity. Whereas macrodiversity describes between-species diversity, microdiversity is the genetic diversity *within* a species or population.

The role microdiversity plays in the eco-evolutionary dynamics of phages cannot be uncoupled from those dynamics of the host system [11, 12, 13]. Bacteriophages and bacteria are involved in a complex coevolutionary arms race between infecting and preventing infection. Microdiversity, both in viruses and their hosts, is essential in maintaining the possibility for long-term host-virus coexistence [14]. Besides host-virus interactions, microdiversity is also necessary to provide ample variation for adaptation upon changing environmental conditions [15, 16]. Microdiversity in marine phages has been shown to be influenced by host availability, environmental stability and virus population dynamics [17, 1]. Unfortunately, despite its importance, microdiversity in bacteriophages is still understudied compared to microdiversity in bacteria [5].

The interplay between host and phage gains an extra dimension when the lysogenic lifestyle of temperate phages is considered in contrast to the lytic lifestyle of virulent (or obligatory lytic) phages [18, 19]. Both virulent and temperate phages can perform the lytic lifecycle: the phage infects the cell, starts replication of its genome and production of necessary structural proteins and finally induces cell lysis to release new virions. Genome replication during the lytic life cycle is affected by the viral mutation rate, which is estimated to be up to two orders of magnitude higher than the mutation rate of the host [20, 21, 22]. The lysogenic lifecycle of temperate phages involves integration of the viral genome into the host genome, instead of immediate production of virions and lysis. This turns the phage into a prophage and the host into a lysogen. The prophage can multiply with the lysogen when the cell divides, or excise itself from the host genome and return to the lytic lifestyle. Depending on biotic and abiotic factors, temperate and virulent phages can be under very different selection pressures. For example, some bacteria can transform into endospores, potentially protecting the prophage from extreme environmental conditions [23]. Furthermore, prophages can exchange genetic material with superinfecting phages or other prophages [24, 11]. However, how the lifestyle of bacteriophages affects their genetic microdiversity is currently unknown.

The ratio of phage lifestyles differs by environment, but even within environments factors like depth can make a difference [25, 17]. Overall estimates indicate a higher degree of lysogeny in soil environments versus marine environments. Besides this lifestyle ratio, soil viruses are incredibly diverse, both in structure and diversity, while also having a high potential impact on soil health through nutrient cycling and microbiome composition [26, 27, 28].

Here we studied the association between phage lifestyle and microdiversity in *Caudoviricetes*, a well-studied class of double-stranded DNA viruses that can have either a virulent or temperate lifestyle [6]. 12 soil-borne bacteriophage datasets, selected for their recent publication and FAIR data (i.e. data was Findable, Accessible, Interoperable and Reusable), were re-analysed. The 12 datasets together contained 537 397 viral operational taxonomic units (vOTUs, approximately species rank viral genomes), which were filtered and analysed according to best current practices [29] using a high-throughput analysis pipeline. Finally, the association between lifestyle (temperate or virulent) and microdiversity was tested on 43 171 vOTUs.

## 2. Experimental procedures

### 2.1. Data collection

Datasets were selected from metagenome studies of soil-based bacteriophages catalogued in the NCBI PubMed database. Additional requirements were recent publishing (*<* 5 years, from December 2025) and data availability of raw reads and vOTUs. Using these criteria, 17 studies were selected for further analysis (Table 1). The vOTUs and raw reads were retrieved for these datasets. vOTUs were downloaded from their respective data repositories. Raw reads were retrieved from SRA by their BioProject ID (Table 1 for IDs) using the NCBI SRA-toolkit (github.com/ncbi/sra-tools) [30]. There were slight differences how the original studies computed the vOTUs from assembled contigs. 14 studies followed the predominant standard for clustering of contigs into vOTUs: at least 95% average nucleotide identity over 85% alignment fraction [1, 31], whereas 3 studies used different parameters. Given the vOTUs from the original studies were used and no *de novo* assembly and phage identification was performed, duplicates may exist between datasets. Hence, all datasets were also reclustered together using PSI-CD-HIT (v4.8.1, using the flags for alignment thresholds “-c 0.95 -G 1 -g 1 -aS 0.85 -aL 0.85”) and the standard definition for vOTUs (at least 95% average nucleotide identity over 85% alignment fraction) to check for redundancy caused by different vOTU clustering definitions between datasets [32]. This re-clustering only resulted in a very small reduction in vOTUs (see Supplemental Information, “Re-clustering of vOTUs using standard thresholds”); thus, we continued with the original vOTUs from each dataset.

**Table 1.** Overview of the 17 datasets used in this study. All datasets are of soil-based viruses, most of them focusing specifically on bacteriophages. Studies that performed size fractioning (*<* 0.22*µ*m) in their sample treatment show the number of viromes (v) or metagenomes (m). If a study had both viromes and metagenomes available, only the number of viromes is shown. Standard clustering parameters are *>*= 95% ANI over *>*= 85% AF. **ANI** = Average Nucleotide Identity, **AF** = Alignment Fraction.

| Dataset ID | Source | BioProject ID | Clustering | # of libraries | # of vOTUs |
| --- | --- | --- | --- | --- | --- |
| AgriSoil | [7] | PRJNA646773 | standard | 15 (v) | 4 065 |
| BodegaBay | [33] | PRJNA913601 | standard | 63 (v) | 12 689 |
| Conifer | [34] | PRJNA1113186 | standard | 90 (v) | 272 017 |
| Glacial | [9] | PRJNA882246 | $\geq 95\%$ ANI over $\geq 80\%$ AF | 24 (v) | 59 685 |
| Hopland | [35] | PRJNA818793 | standard | 44 (v) | 6 088 |
| Intertidal | [36] | PRJNA957716 | standard | 96 (m) | 20 102 |
| Land-use | [37] | PRJNA691683 | standard | 50 (v) | 61 106 |
| MediGrass | [38] | PRJNA1264504 | standard | 59 (v) | 22 725 |
| NCalHabitat | [39] | PRJNA831438 | standard | 30 (v) | 3 432 |
| NewMexico | [40] | PRJNA1263537 | standard | 11 (v) | 3 233 |
| PermaThaw | [41] | PRJNA888099 | standard | 20 (v) | 9 963 |
| Rhizo | [42] | PRJNA935076 | $\geq 95\%$ ANI over $\geq 80\%$ AF | 12 (v) | 13 335 |
| Spruce | [43] | PRJNA666221 | standard | 10 (v) | 5 006 |
| UKagri | [44] | PRJEB49313 | $\geq 95\%$ ANI over $\geq 95\%$ AF | 15 (v) | 1 059 |
| Watershed | [45] | PRJNA1168581 | standard | 48 (m) | 2 743 |
| WetUp | [46] | PRJNA859194 | standard | 152 (v) | 12 259 |
| Wildfire | [47] | PRJNA1147726 | standard | 39 (v) | 70 135 |

### 2.2. vOTU quality assessment and mapping of reads

The files containing the vOTUs (for each dataset) were used as supplied by the authors of the respective publications. All vOTUs were analysed with CheckV (v1.0.3, “end_to_end” pipeline, database version 1.5) and filtered on at least 80% estimated genome completeness [48].

Raw reads of each library were mapped against the vOTUs of the respective study with Bowtie2 (v2.4.4), using the “very-sensitive” setting [49]. Unmapped reads were discarded during the mapping process, using the Bowtie2 “–no-unal” flag. Bowtie2 SAM output files were converted to BAM files, sorted using SAMtools (v1.21) and merged within each dataset [50]. Coverage of contigs by mapped reads was calculated using CoverM (v0.7.0, using the contig option, “mean” coverage calculation method) [51]. vOTUs were filtered for a minimum coverage of 10x and breadth of coverage of at least 80%. These thresholds were necessary for downstream micrdiversity analysis, primarily ensuring accurate (high coverage) microdiversity estimation of the full genome (high breadth). All vOTUs were also assigned taxonomic information using geNomad (v1.11.1, default settings)[52] and filtered for being classified as *Caudoviricetes*, as the lifestyle prediction tools were designed for *Caudoviricetes* viruses [53, 54].

### 2.3. Prediction of vOTU lifestyle

The lifestyle of all vOTUs was predicted using PhaTYP (as included in PhaBOX2 v2.1.11) [53], PhaStyle (v0.1) [54] and verified by checking for integrases with DIAMOND (v2.0.8) [55]. PhaStyle uses a ProkBERT genomic language model to detect similarity of query sequences to phages with known lifestyles. It uses a binary classification model: the negative class in the model is the temperate lifestyle, the positive being virulent. PhaStyle was run using a GPU, greatly accelerating the processing speed compared to CPU-based usage. PhaStyle was run using the “neuralbioinfo/PhaStyle-mini” model available online (huggingface.co/neuralbioinfo/prokbert-mini). PhaTYP also uses a BERT-based model, but differs from PhaStyle in the method of embedding and masking sequences [56, 54]. These tools do not identify other lifestyles, such as pseudolysogeny or chronic phages.

vOTUs were also checked for presence of lysogeny marker genes using the cluster of orthologous genes COG0582, COG2915, COG3311, COG4570 and COG4974 from the COG database and DIAMOND (sensitive preset, e-value *<*1e-20 and minimum query coverage per High-Scoring Pair of 80%)[55, 57]. These COGs were chosen for their role in the lysogenic lifecycle, specifically integration (COG0582, COG4974) and lysogen/prophage regulation (COG2915, COG3311, COG4570). The COGs together consisted of 53 451 protein sequences and formed the database against which the proteins from all vOTUs, as predicted by Prodigal, were queried [58]. The final lifestyle classification was determined by the overlap between PhaTYP and PhaStyle and presence of a DIAMOND hit with a lysogeny marker protein sequence for the temperate lifestyle or absence of such a hit for the virulent lifestyle.

### 2.4. Microdiversity analysis of filtered vOTUs

All vOTUs with mapped reads were used as input for calculating microdiversity using inStrain (v1.9.1) [59]. inStrain applies a 95% ANI filter for each aligned reads, reducing spurious mappings. Default parameters were used for variant calling within inStrain (SNP coverage *>*= 5, SNP frequency *>* 5%, FDR = 1e-06). The main output metric of interest was the nucleotide diversity (*π*) per vOTU, included in the scaffold output files. Additionally, pN/pS ratios in predicted gene sequences were computed by inStrain and retrieved from the gene output files. pN/pS is the ratio between non-synonymous and synonymous mutations in gene coding sequences and can be an indication of selective pressure on a genetic sequence. For datasets with at least 10 vOTUs per lifestyle, the ratio between single library microdiversity and global microdiversity (i.e. the per-dataset approach) was also calculated.

### 2.5. Association analysis of lifestyle-microdiversity relationship

The association analysis between phage lifestyle and microdiversity was performed using R (v4.4.2) [60] and additional libraries from the tidyverse were used for data handling and visualisation [61]. We required there to be at least 10 vOTUs in each lifestyle group per dataset analysed. A Wilcoxon rank-sum test was used to calculate the statistical relationship between the two lifestyle groups and their respective microdiversity.

### 2.6. Functional annotation of viral genes

vOTUs belonging to the Land-use dataset also underwent functional prediction of genes. For this, genes were first predicted using Prodigal (v2.6.3, anonymous mode) [58]. After this, eggNOG mapper (v2.1.13, default settings) was used in combination with the eggNOG v5 database to predict the function of genes in vOTUs [62, 63]. Annotations were parsed for keywords in the GFF fields for predicted PFAM domains (“em_PFAMS”) and description (“em_desc”). Since the focus was on essential genes, only the genes containing the following values in the GFF fields “em_desc” amd “em_PFAMs” were highlighted during visualisation: “terminase”, “capsid / head / coat”, “portal”, “tail”, “helicase”, “integrase” and “recombinase”.

### 2.7. SRR2strain

The workflow presented here has been largely integrated into a single Snakemake pipeline (v9.3.3), called SRR2strain (https://github.com/thattimetho/SRR2strain, Supplemental Figure 1). Snakemake is a python-based workflow management system, that allows the linking of input and output of tools to automate workflows [64]. The steps mentioned here, from the retrieval of the raw reads up until the microdiversity analysis using inStrain, have been incorporated into the pipeline. The acquisition of the respective vOTU databases, functional annotation of Land-use vOTUs and final analysis using R scripts are not incorporated in the pipeline.

## 3. Results

### 3.1. Viromes facilitate stringent filtering required for reliable microdiversity estimation

A systematic literature search was performed for studies on bacteriophages and soil environments, taking into account the recency of publication and availability of data. This search resulted in 17 datasets. These 17 datasets formed the starting point of our analysis into the relationship between phage lifestyle and microdiversity. Of the 17 datasets, two contained only metagenomes, while the rest contained only viromes or both viromes and metagenomes (Table 1). The datasets are from diverse soil environments (e.g. coastal, agricultural, urban). These datasets were used as input for the SRR2strain pipeline (Supplemental Figure 1, also see subsection 2.7), which predicts phage lifestyle and estimates phage microdiversity. As part of this pipeline, sequenced reads are mapped back onto the vOTUs to allow for the calculation of microdiversity downstream (using inStrain) [59]. During this mapping process, we observed that the metagenomes showed extremely poor mapping efficiency (less than 3% mapped reads, Supplemental Table 1). This was irreconcilable with the thresholds on minimum mapping breadth and coverage necessary for accurate microdiversity estimation. For this reason, the samples with only metagenomes (Watershed and Intertidal) were excluded from the analysis. Only viromes were considered for the remaining datasets and analyses, which led to a significant improvement in mapping efficiency (see Supplemental Table 1).

The 15 remaining datasets were filtered for sufficient mapping coverage and breadth, and vOTU completeness. This led to the removal of Rhizo, Spruce and UKagri, as these datasets did not have sufficient vOTUs (at least 10 vOTUs per lifestyle, see methods) left for lifestyle microdiversity association analysis. From the other 12 datasets, 117 022 out of 537 397 vOTUs remained after these (mapping) quality control measures were applied. Subsequent taxonomic classification was performed using geNomad and resulted in successful classification of 116 445 vOTUs (99.5% of 117 022 total). Of these, 115 922 (99.6%) were predicted to belong to the *Caudoviricetes* class of viruses. Of the remaining non-*Caudoviricetes* 523 vOTUs, 411 were assigned as *Tectiviridae*, 83 as *Priklausovirales*, with the remainder spread across various *Monodnaviria* and *Varidnaviria* subtaxa. For subsequent analyses and results, only the vOTUs classified as belonging to the *Caudoviricetes* class were considered.

### 3.2. Virulent phages comprise majority in lifestyle distribution

Phage lifestyle, either virulent or temperate, was predicted for the 115 922 *Caudoviricetes* vOTUs after quality filtering using a combination of tools (PhaTYP, PhaStyle and DIAMOND)(Table 2). This resulted in 43 171 vOTUs which were assigned the same lifestyle by both PhaTYP and PhaStyle, requiring temperate phages to have a lysogeny marker protein detected by DIAMOND. Of these 43 171 vOTUs, 11 524 (27%) were classified as having a temperate lifestyle and 31 647 (73%) as having a virulent lifestyle. The ratio between virulent and temperate phages per study was similar between all 12 studies (Figure 1). The lifestyle ratio changed slightly when corrected for relative abundance of vOTUs, leading to temperate phages increasing in proportion (Supplemental Figure 2). 72 751 vOTUs that passed the initial stringent quality checks were discarded due to disagreement between lifestyle prediction methods (Figure 1A).

**Fig. 1.**
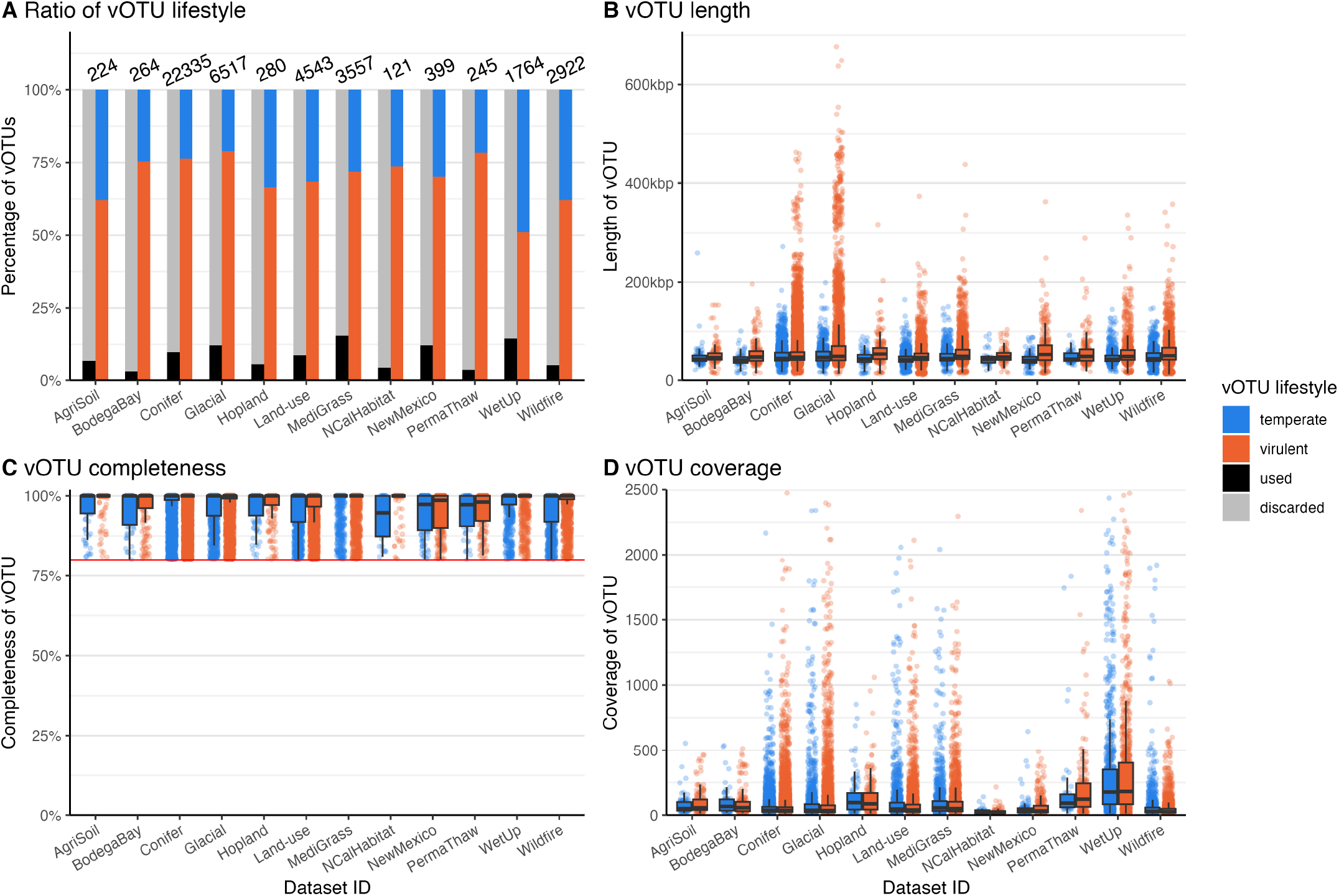
Overview of vOTU related statistics. (**A**). Ratios of total and used data, and the ratio between temperate and virulent classified vOTUs. The numbers indicate total vOTUs used for microdiversity analysis. (**B**). Length distribution of used vOTUs among the 12 datasets. Strikingly, only the Glacial and Conifer study show multiple viral genome lengths above 400kbp. (**C**). Predicted completeness of used vOTUs. The used threshold of 80% is indicated with the red line. (**D**). Mapping coverage of used vOTUs. The threshold for coverage was put at 10x.

**Fig. 2.**
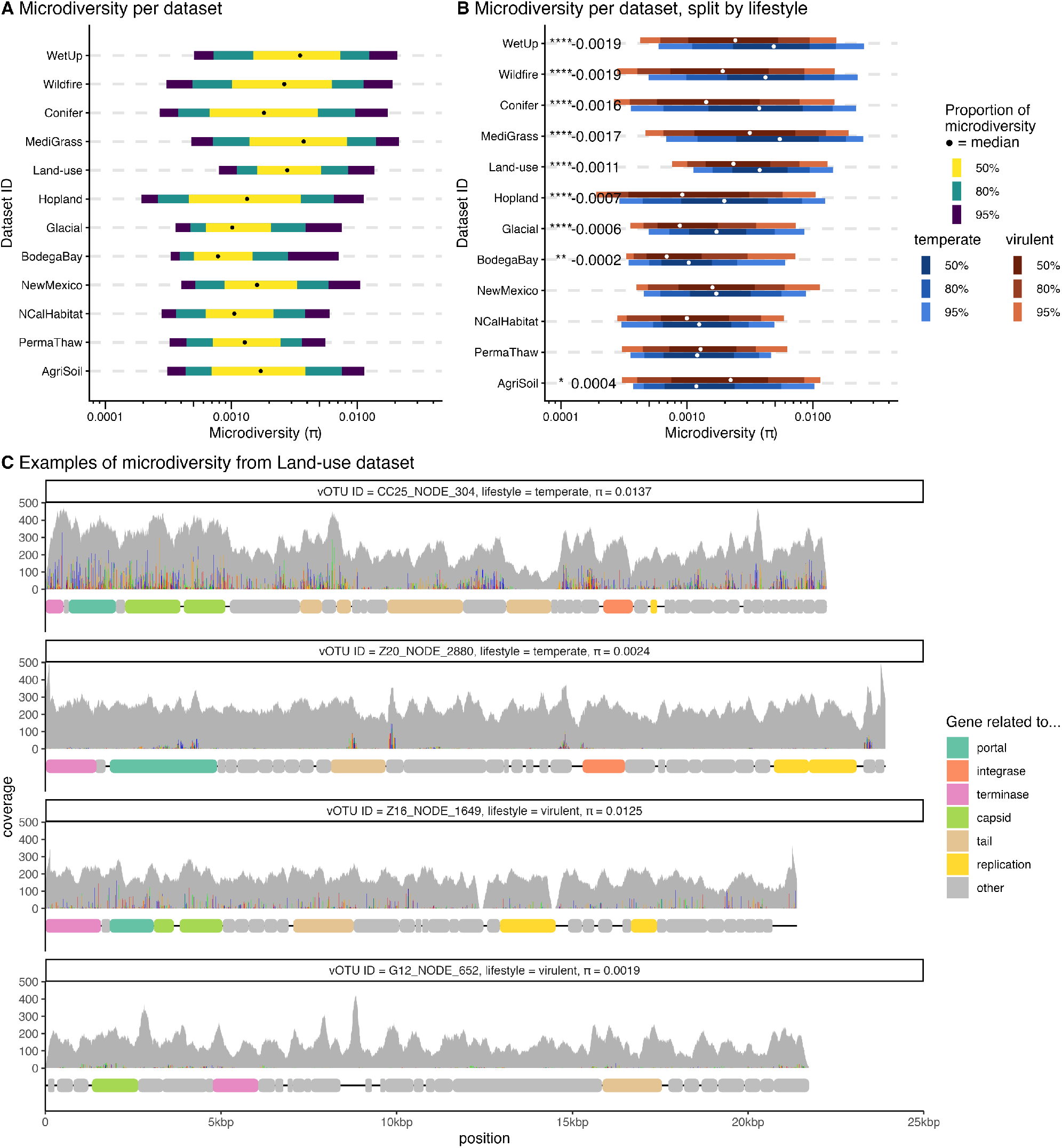
Distribution of microdiversity in temperate phages and virulent phages across 12 datasets. (**A**,**B**) Datasets are sorted by difference in microdiversity between lifestyles. (**A**) Interval plot of microdiversity distribution of vOTUs. (**B**) Interval plot of microdiversity distribution split by lifestyle indicating significance (**** - p*<*0.0001, *** - p*<*0.001, ** - p*<*0.01, * - p*<*0.05, Wilcoxon rank sum test, two sided) and value next to significance indication shows estimated difference between distributions. Higher microdiversity in temperate phages is shown by the more right skewed microdiversity distributions of the temperate phages. (**C**) Four Land-use vOTUs showing different microdiversity and lifestyle. Of each lifestyle, both a high and low microdiversity example is given. Coverage for the dataset (read identity *>*95%) and gene annotation across the genomes are shown.

**Table 2.**
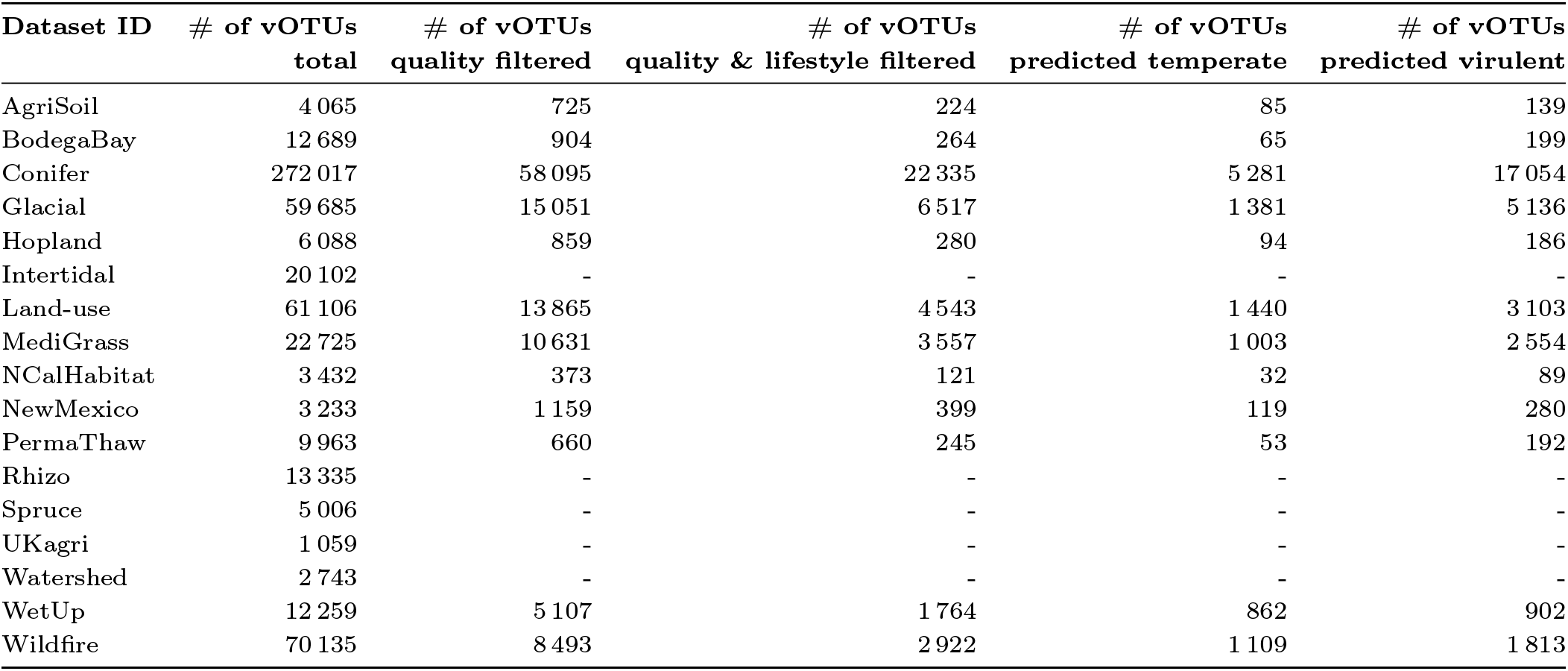
Overview of datasets used as input for microdiversity analysis. The total column indicates input data, whereas the quality filtered column indicates quality and taxonomy filtered data. Taxonomic classification resulted in almost all quality filtered vOTUs being classified as *Caudoviricetes* viruses. The last two columns are based on total number of quality, taxonomy, and lifestyle filtered vOTUs, split by lifestyle. The sum of the lifestyles is shown in the “# of vOTUs quality & lifestyle filtered” column.

| Dataset ID | # of vOTUs<br>total | # of vOTUs<br>quality filtered | # of vOTUs<br>quality & lifestyle filtered | # of vOTUs<br>predicted temperate | # of vOTUs<br>predicted virulent |
| --- | --- | --- | --- | --- | --- |
| AgriSoil | 4 065 | 725 | 224 | 85 | 139 |
| BodegaBay | 12 689 | 904 | 264 | 65 | 199 |
| Conifer | 272 017 | 58 095 | 22 335 | 5 281 | 17 054 |
| Glacial | 59 685 | 15 051 | 6 517 | 1 381 | 5 136 |
| Hopland | 6 088 | 859 | 280 | 94 | 186 |
| Intertidal | 20 102 | - | - | - | - |
| Land-use | 61 106 | 13 865 | 4 543 | 1 440 | 3 103 |
| MediGrass | 22 725 | 10 631 | 3 557 | 1 003 | 2 554 |
| NCalHabitat | 3 432 | 373 | 121 | 32 | 89 |
| NewMexico | 3 233 | 1 159 | 399 | 119 | 280 |
| PermaThaw | 9 963 | 660 | 245 | 53 | 192 |
| Rhizo | 13 335 | - | - | - | - |
| Spruce | 5 006 | - | - | - | - |
| UKagri | 1 059 | - | - | - | - |
| Watershed | 2 743 | - | - | - | - |
| WetUp | 12 259 | 5 107 | 1 764 | 862 | 902 |
| Wildfire | 70 135 | 8 493 | 2 922 | 1 109 | 1 813 |

In order to identify and assess any confounding variables, various other metrics, like mapping breadth, coverage, vOTU length and completeness, have also been tested in relation to each other and to nucleotide diversity (Spearman correlation, alpha = 0.05)(Supplemental Figure 3). The only consistent significant correlation found by this analysis was between mapping coverage and breadth. Some inconsistent correlations have also been found, like those between microdiversity and genome length, and microdiversity and pN/pS. The association of lifestyle with the aforementioned variables has also been tested (Wilcoxon rank-sum test, two-tailed). Some of these metrics show a significant statistical association with vOTU lifestyle, but the estimated true difference between the groups is very small (Figure 1B-C and Supplemental Table 2).

### 3.3. Overview of high quality viral datasets

After filtering on mapping and vOTU quality and taking congruent lifestyle predictions, the final assembled dataset consisted of 12 datasets, containing 43 171 vOTUs (8% of all 537 397 vOTUs in these datasets, see also Table 2). The included vOTUs were not distributed equally among the 12 studies: ~50% originated from the Conifer dataset. On the dataset level, average mapping breadth and estimated vOTU completeness showed similar values among the 12 studies, while average vOTU length and average coverage showed more variability between datasets (Figure 1, Supplemental Table 3 and Table 4). Median mapping coverage showed values between 21x (NCalHabitat) and 186x (WetUp).

When split by lifestyle, correlation analysis (Spearman correlation, alpha = 0.05, see Supplemental Figure 3 and also Supplemental Table 4) indicated a general correlation between microdiversity and length, with eight out of 12 datasets showing a negative correlation. This was also supported by an association analysis (Wilcoxon rank-sum test, two-tailed, see Supplemental Table 2). Since the estimated effect is small and microdiversity is corrected for genome length, no additional corrections were applied as a result of this.

### 3.4. Soil-borne bacteriophage microdiversity shows association with lifestyle

Microdiversity (*π*) present within the vOTUs was calculated to identify and analyse any association with phage lifestyle. Generally, microdiversity levels were similar between studies and consistent with the microdiversity reported in the original studies [37, 9]. Overall, the studies showed quite a wide range of microdiversity (Figure 2A), ranging from a median of 0.0010 (BodegaBay) to a median of 0.0054 (MediGrass). When comparing single libraries to the global (pooled) dataset, microdiversity per library is generally more than half of the dataset microdiversity (see Supplemental Table 4).

We find that nine out of 12 datasets showed a significant difference in the level of microdiversity detected for each lifestyle (Figure 2B and Supplemental Table 5). Only one of these (AgriSoil) shows a higler level of microdiversity in virulent phage. In contrast, the vast majority of the significant datasets showed a higher level of microdiversity in temperate phages compared to virulent phages, where the WetUp dataset showed the most distinct microdiversity distributions between the two lifestyles (p = 6.4e-36, estimate = 0.0019, Wilcoxon rank-sum test, two-tailed). The Conifer dataset contained the most vOTUs out of all datasets, which influenced the statistical significance of the difference in microdiversity between the lifestyles (p = 0, estimate = 0.0016, Wilcoxon rank-sum test, two-tailed). Overall, the association between lifestyle and microdiversity was robust when using different lifestyle prediction methods and when correcting for skewed lifestyle ratios (see Supplemental Table 5 and Supplemental Figure 4).

When inspecting read mapping across example genomes, we find reasonable coverage patterns, containing no clearly depleted or significantly enriched areas of the genome (Figure 2C). The coverage of the vOTUs also does not seem to correlate strongly with the amount of microdiversity, which is supported by the general correlation analysis on the Land-use dataset (Spearman correlation, alpha = 0.05, Supplemental Figure 3). The localisation of microdiversity seems to vary by vOTU, with Z20_NODE_2880 seemingly showing more microdiversity in the terminase and portal protein predicted proteins compared to the rest of the genome. Meanwhile, Z16_NODE_1649 shows a more even distribution of microdiversity throughout the vOTU.

In addition to vOTU-wide microdiversity analysis, the effect of lifestyle on gene-level characteristics was also studied. Besides general statistics, like length and breadth (see Supplemental Figure 5),

To analyse the association of selective pressure on genes with lifestyle, we analysed the pN/pS ratios as calculated by inStrain. Generally, ratios between 0 and 1 indicate purifying selection, ratios around 1 are a sign of neutral protein evolution and ratios above 1 indicate positive selection. We find that the majority of pN/pS ratios is below 1, indicating purifying selection on the vOTUs (Figure 3). Phage lifestyle showed a significant association with pN/pS in 11 out of 12 datasets (Wilcoxon rank sum test, see also Supplement Table 6), where temperate phages showed higher pN/pS than virulent phages. Lastly, correlation analysis (Spearman correlation, alpha = 0.05) shows a strong positive correlation between the pN/pS ratio and microdiversity in 10 out of 12 datasets (Supplemental Figure 3 and Supplemental Figure 6).

**Fig. 3.**
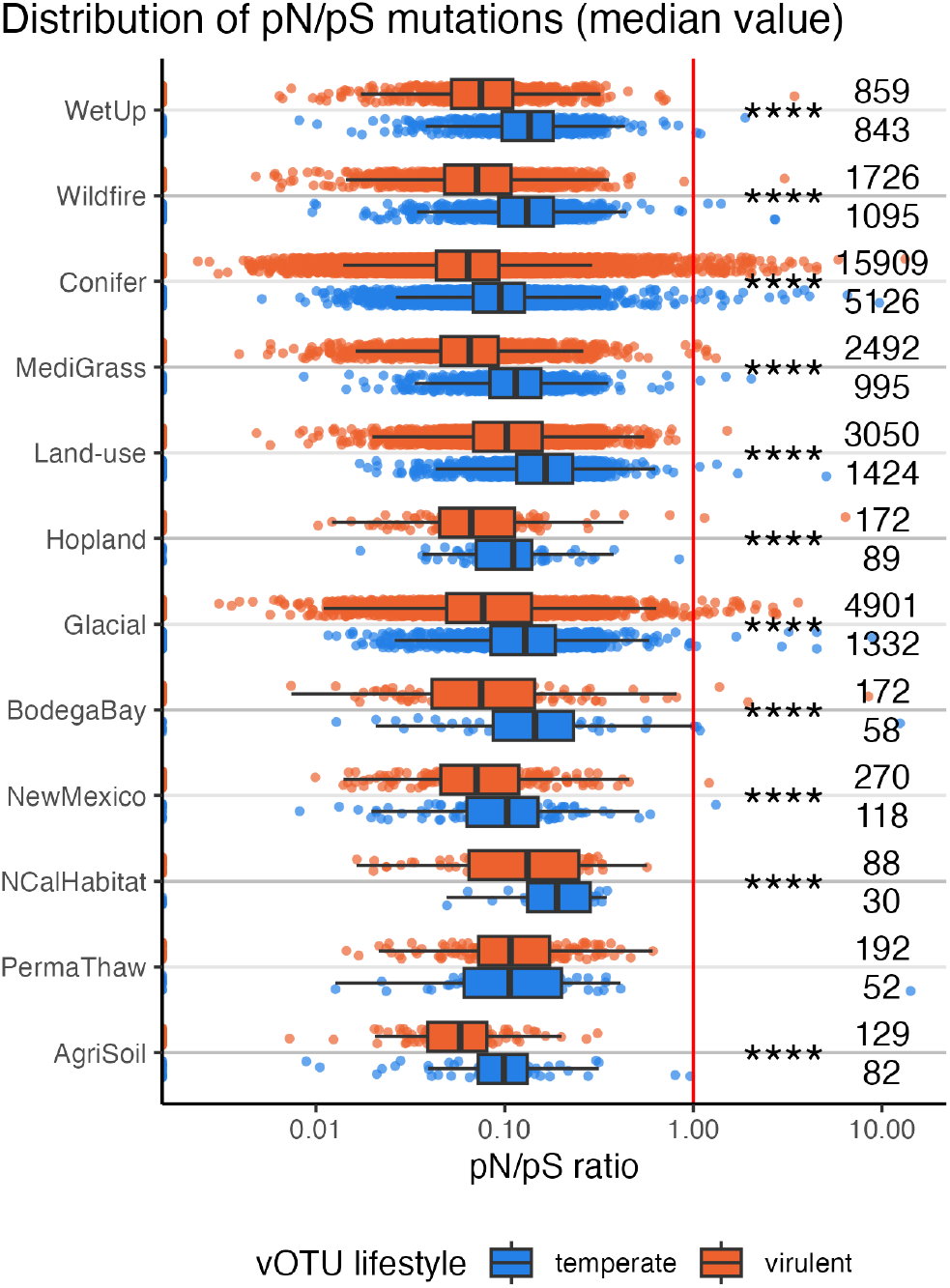
pN/pS ratios per dataset split by lifestyle. Temperate phages show consistently and significantly higher pN/pS ratios than virulent phages. Difference in pN/pS between lifestyle was statistically tested with a Wilcoxon rank sum test, resulting in significant differences in the pN/pS values of 11 of 12 datasets between temperate and virulent lifestyles as indicated by the asterisks (see Supplemental Table 6). The numbers on the right indicate the number of vOTUs that were included in the Wilcoxon rank sum test and plot.

## 4. Discussion

Advances in high-throughput, cost-effective sequencing and improved assembly have led to an expanding set of highquality DNA viromic datasets from diverse environments, including soils. Leveraging these resources, we find that phage lifestyle is associated with microdiversity across different soil datasets. Using stringent filters for both read mapping and lifestyle prediction, we obtained confident association statistics for 12 datasets; nine of the 12 showed significant lifestyle-associated differences, with eight showing temperate phages displaying higher microdiversity (Figure 2). Across those eight datasets, statistics show an estimated ~36% difference in microdiversity between the two lifestyles. Overall microdiversity levels and lifestyle distribution were in line with those reported by previous soil-focused studies [37, 36, 9]. Furthermore, the pN/pS ratio indicates of the type of selection pressure on protein-coding genes. We find generally low pN/pS ratios, indicating that the vOTUs are under purifying selective pressure and pN/pS rations are generally lower for virulent phages (Figure 3). Purifying selection can be a sign of a relatively unperturbed environment, thus not favouring new variants or gene functions to allow for new niche occupation [65]. However, soil is usually seen as a very dynamic environment, especially topsoil, which is generally sampled [43, 66]. Notably, genome-wide purifying selection might hide the signatures of positive selection on only a few protein regions. Additional analyses are necessary to investigate how the distribution of these mostly synonymous mutations is spread across the gene repertoire of phages and whether these mutations localise in certain groups of genes.

As this project aimed to connect microdiversity with phage lifestyle, accurate microdiversity estimation is key. Micro-diversity analysis requires a high read depth, as to filter out sequencing errors and provide certainty on the amount of genetic diversity present. Our initial collection of datasets contained two studies with metagenomes only rather than (size-fractionated) viromes. However, mapping the metagenomic reads against the vOTUs resulted in low mapping efficiency and low coverages of vOTUs and consequently to a low number of vOTUs with sufficient coverage. This is consistent with previous reports that metagenomes do not contain viral reads at sufficient coverage for downstream analysis [7, 67]. Soil samples are especially hard to sequence for viruses using metagenomics, underlining the need for viromes further [68, 67]. Even when using viromes, the strict quality thresholds we imposed substantially reduced analysable vOTUs. For example, in the BodegaBay dataset, this led to a reduction in vOTUs of 98%, of which: 64% did not pass the quality thresholds (but could be assigned a lifestyle with certainty), and 34% lacked confident lifestyle assignment. For three datasets (Rhizo, Spruce, and UKagri) the stringent filtering resulted in too few vOTUs for association analysis, which could be due to the low number of libraries and total reads in these datasets (Table 1, Supplemental Table 1). This results in a low overall coverage and highlights the importance of deeply-sequenced viromes for such fine-grained analyses. The pooling of viromes, as done in this study, is a way to further increase the number of vOTUs with high coverage and breadth, but does mean the analysis and results should be considered to be on an environmental level, not a local one. This is also seen when comparing per-sample microdiversity and pooled microdiversity, with all datasets showing a higher pooled microdiversity (Supplemental Figure 7).

Not only the data used influences the output achieved, so do the various clustering and mapping methods, tool settings and filtering steps. For example, the pipeline only includes a single mapping tool, Bowtie2. Different mappers with different settings could influence the mapping efficiency and accuracy. Nevertheless, we expect only a limited effect of using another mapping tool, given that the reads from all datasets were mapped with the same tool and the mapping is independent of phage lifestyle. The subsequent filtering using both coverage and breadth was aimed at reducing spurious mappings. In addition to these thresholds, inStrain was run using default filtering steps, which also includes an ANI cutoff of 95% of mapped reads. Overall, given the many strict thresholds set on vOTU and mapping quality, we are confident that the results presented here are a realistic representation of the microdiversity present in the respective datasets.

In this study, bacteriophage lifestyle is hypothesized to be an explaining variable of microdiversity, and as such, lifestyle prediction needs to be as accurate and precise as possible. Because of this, we combined two sequence-based lifestyle prediction tools, namely PhaTYP and PhaStyle, with DIAMOND search against lysogeny marker genes [55, 56, 54]. In addition to choosing the overlap between these three tools, various combinations of tools have also been tested, which shows that the microdiversity results are robust towards the chosen lifestyle prediction tool (Supplemental Table 5). PhaTYP and PhaStyle were chosen for their recent development and extensive benchmarking against established but older tools [56, 54]. Despite similar conceptual underpinnings, PhaTYP and PhaStyle yielded substantial variance in lifestyle calls (Supplemental Figure 8). This subsequently led to 72 751 vOTUs being filtered out due to disagreement between the lifestyle prediction results (Table 2). This difference between tools is the exact reason we combine the predictions of multiple tools. It also highlights the difficulty of accurate lifestyle prediction. The actual ratio between virulent and temperate phages in soil may differ from our current estimates from the filtered datasets, either due to lifestyle prediction methods or sampling bias. To improve lifestyle prediction *a posteriori* for studies containing both metagenomes and viromes, assembled bacterial genomes could be checked for prophages also present in the viral fraction, thus indicating active lysogeny. Other options of integrating orthogonal evidence to reduce uncertainty in lifestyle assignment include experimental methods like studying the inducible fraction or bioinformatic methods like prophage signature analysis or long-read phage-host linkage [25, 69, 26, 70].

Our results indicate a consistent difference in pN/pS ratio between lifestyles and indicate an impact of lifestyle on microdiversity. In order to explain these differences, we can take a look at the various evolutionary factors that influence viral populations. Selective sweeps and mutation rates are expected to contribute to microdiversity particularly in large, rapidly reproducing viral populations. The exact link between these processes and the consistent lifestyle-dependent differences we detect is not clear. Several non-mutually exclusive hypotheses could account for elevated micro-diversity in temperate phages. First, temperate phages can alternate between lysogenic and lytic replication, potentially increasing opportunities for accumulation and maintenance of within-species polymorphisms across fluctuating ecological conditions and host states [71, 23, 72, 73]. These fluctuating ecological conditions are expected to lead to strong population bottlenecks, especially for virulent phages, as those do not have the option to integrate themselves into a host besides pseudolysogeny. Pseudolysogeny is a stalled state of a phage infection, and could mean virulent phages behave similar to the prophage state of temperate phages [74]. Nonetheless, the harsh world that phages live in is expected to lead to hard selective sweeps, where a genome carrying advantageous mutations increases in frequency leading to a substantial reduction in diversity [75]. Due to large phage population sizes, we expect even a minor advantageous mutation to rapidly fix in this environment. These hard selective sweeps may remove a part of the microdiversity from the virulent phage population, but not per se from temperate phages which are integrated into a host. This could explain both pN/pS lifestyle association, as well as the correlation between microdiversity and pN/pS, either by temperate phages “hiding” from natural selection, or virulent phages losing diversity through background selection and population bottlenecks [19]. Second, integration into diverse host genomes may expose temperate phages to host-mediated selection and recombination landscapes, which can elevate microdiversity relative to strictly lytic counterparts [76, 77, 78]. Lastly, recurrent induction events may repeatedly reseed the temperate phage population with genetically diverse lineages that have diversified due to genetic drift and mutation during prolonged periods of integration [79, 19, 23].

Soil environments are highly spatially heterogeneous, leading to microcosms containing spatially structured bacterial and bacteriophage populations. Prior work has shown that soil viral communities can harbour substantially greater micro-diversity than those from marine systems, likely due to this spatial heterogeneity [1, 39, 37]. The unique spatial properties of soil might also be essential for the pronounced lifestyle signal we observe, given that spatial heterogeneity can greatly influence both phage and host population dynamics [80, 81, 16, 39, 35]. Moreover, while the initial systematic dataset search did not take into account the diversity of sampled biomes, it is worth mentioning that the identified lifestyle-microdiversity association is present across multiple different biomes, like the forests of the Conifer and Wildfire study, and the multiple grassland biomes of the Hopland and MediGrass studies [35, 34, 47, 38]. Yet, while the lifestyle-microdiversity association seems robust, there is one dataset, AgriSoil (Figure 2), showing the opposite trend to the other eight significant associations. However, this dataset has relatively low read numbers and the bootstrapping shows less consistency than for the other significant associations (Supplemental Table 1 and Supplemental Figure 4). Thus, this difference might be caused by methodological differences to the other datasets or it could be a genuine biological result, where specific ecological factors in the environment contribute to the higher microdiversity in virulent phages or the lower microdiversity in temperate phages.

In sum, across multiple high-quality soil viromics datasets, phage lifestyle impacts microdiversity, generally leading to higher microdiversity in temperate phages than virulent phages, a pattern robust to alternative prediction tools and filtering thresholds and present throughout biomes. This suggests that lifestyle-linked ecological and evolutionary processes contribute to the association between lifestyle and micro-diversity. Limited by the data and methods available, we can only hypothesize on potential explanations for this apparent association between lifestyle and microdiversity at the moment. Further investigation, taking into account the complexity of soil environments and the location of diversity across different genomic regions, is warranted to identify the ecological and evolutionary mechanisms driving the consistent lifestyle-microdiversity association.

## 5. Author contributions

TB, HD and AK conceptualization. TB data curation. TB formal analysis. HD and AK funding acquisition. TB investigation. TB and AK methodology. TB, HD and AK project administration. TB software. HD and AK supervision. TB visualization. TB writing - original draft. TB, HD and AK writing - review and editing.

6. Acknowledgements

We would like to thank Marnix Medema and Bas Zwaan for their guidance during the design and final stages of this project. Furthermore, we would like to mention Chen Chen, Dimitris Karapliafis, Laura Patino Medina and Sanne Keizer for their comments on the manuscript.

## 8. Data Availability

A list of IDs and locations for raw read data and vOTU datasets as well as data generated and used for generating visualisations in this study has been uploaded to Zenodo (https://doi.org/10.5281/zenodo.17988606). Raw read data of publications used in this study are available on the NCBI SRA database. vOTU datasets of publications used in this study were retrieved from their respective online data repositories.

## 9. Code Availability

The SRR2strain snakemake pipeline and all other scripts/code generated for this study are available on GitHub (https://github.com/thattimetho/SRR2strain). This repository also contains information on all the tools and libraries used in the processing or visualisation of data.

