## Supplemental Figures and Tables for "Temperate versus virulent phage lifestyle impacts microdiversity in soil environments"

### Supplementary Information and Figures - Microdiversity is higher in temperate than in virulent bacteriophages from soil environments

**Thomas E.P. de Bruijn, Hilje M. Doekes, Anne Kupczok**

#### Re-clustering of vOTUs using standard thresholds

Re-clustering all the 579,642 vOTUs from the 17 virome datasets using a single set of thresholds (95% average nucleotide identity over at least 85% of the aligned fraction) resulted in 576,980 vOTU clusters, of which 574,460 were singletons (99.6%). Of the 2,520 non-singleton clusters, 2,409 contained two vOTUs, 96 contained three vOTUs, 13 contained four vOTUs, and two clusters contained five and eight vOTUs respectively. There was minor clustering over datasets, exclusively in clusters consisting of two vOTUs. Clusters containing more than two vOTUs only showed clustering within datasets. With the vast majority of the 579,642 vOTUs being clustered into singletons, the unclustered dataset was used for further analyses.

#### Supplemental Figures

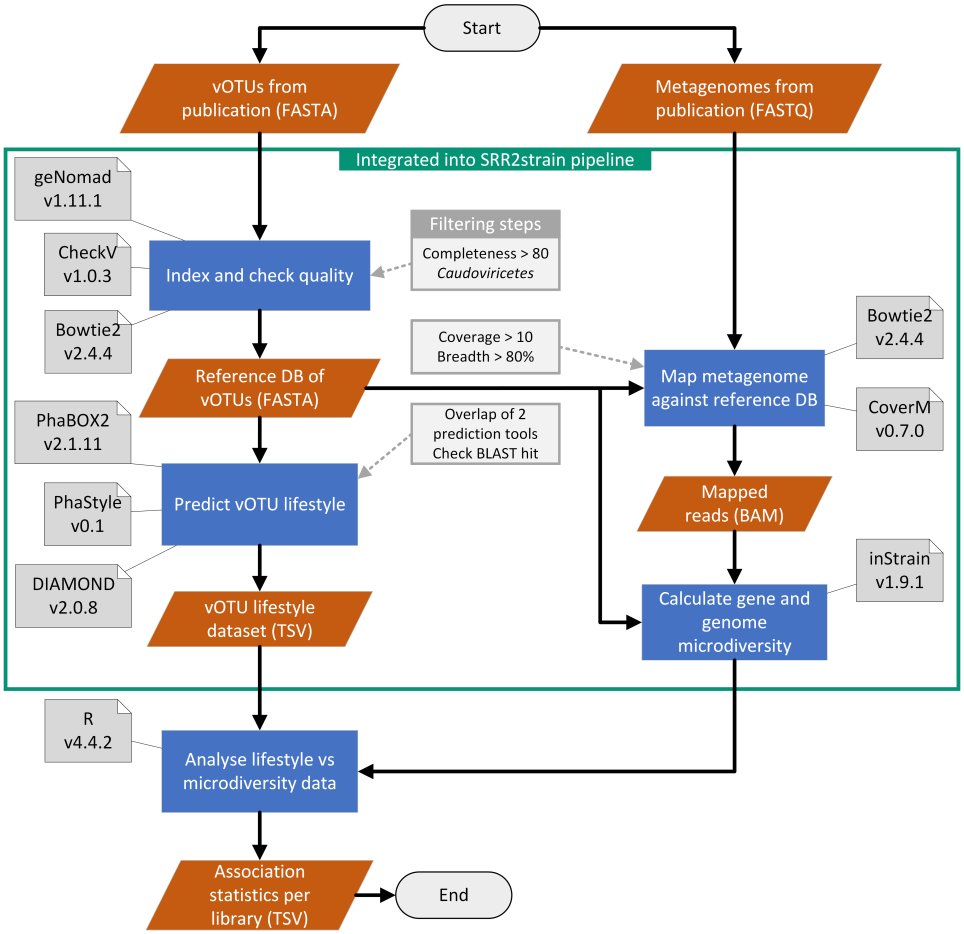


**Supplemental Figure 1. Flowchart showing the steps taken and tools used in this study** to process raw reads and vOTUs resulting in microdiversity and lifestyle association information. A large part of the steps shown here has been integrated into a snakemake pipeline, enabling high reproducibility and transparency. Version numbers shown next to the tools above can also be found in the Conda environment .yml files. The pipeline is meant to run on a per-dataset basis.

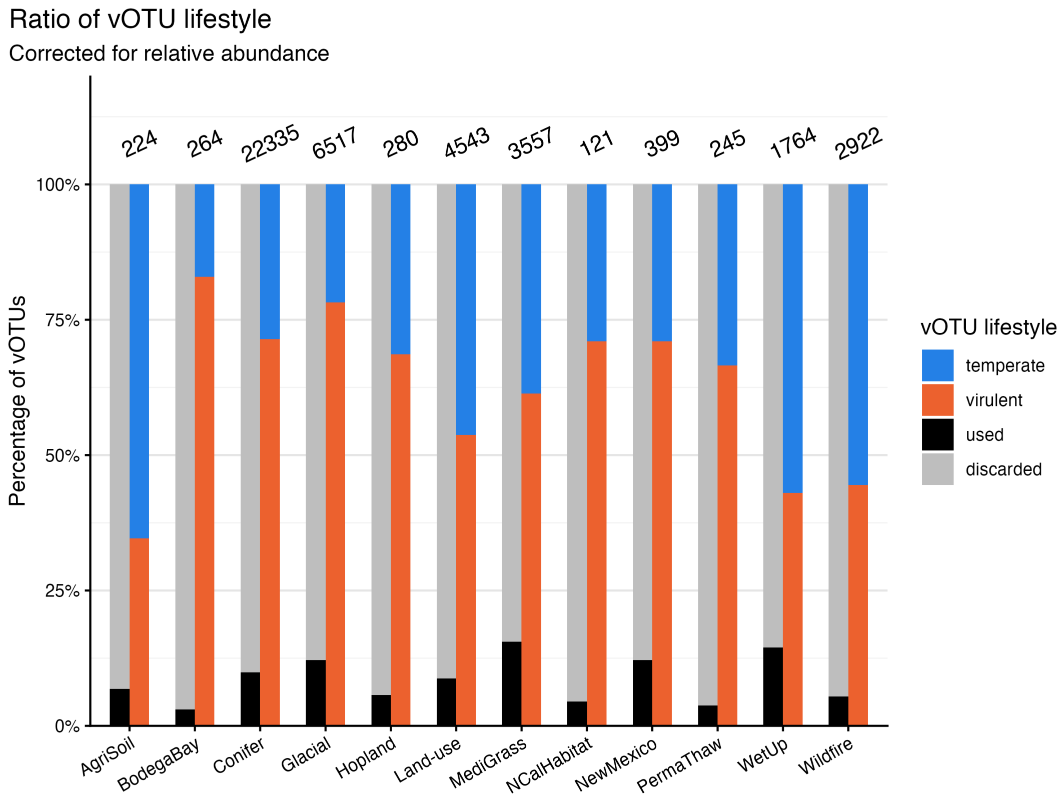


**Supplemental Figure 2. Overview of number of vOTUs used in analysis, corrected using relative abundance**. Ratios of total and used data, and the ratio between temperate and virulent classified vOTUs. The numbers indicate discarded vOTUs due to quality issues and total vOTUs used for microdiversity analysis. Bar charts used relative abundance as weights, thereby correcting for relative abundance.

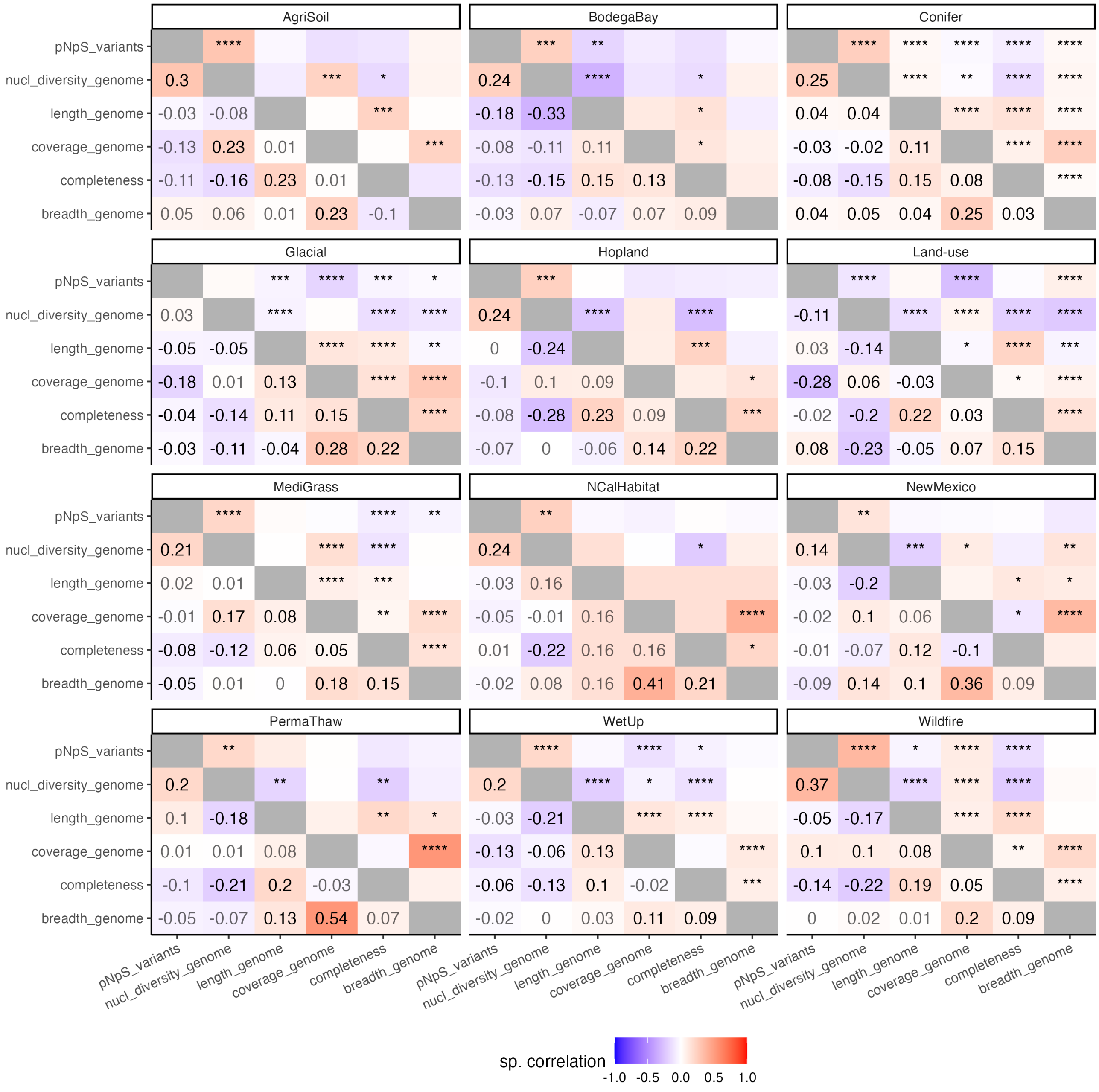

**Supplemental Figure 3. Spearman correlations of various quality metrics and nucleotide diversity (i.e. microdiversity) per dataset.** Correlations were calculated of the twelve datasets using a Spearman correlation test. Correlation coefficient is shown for p-value < 0.05.

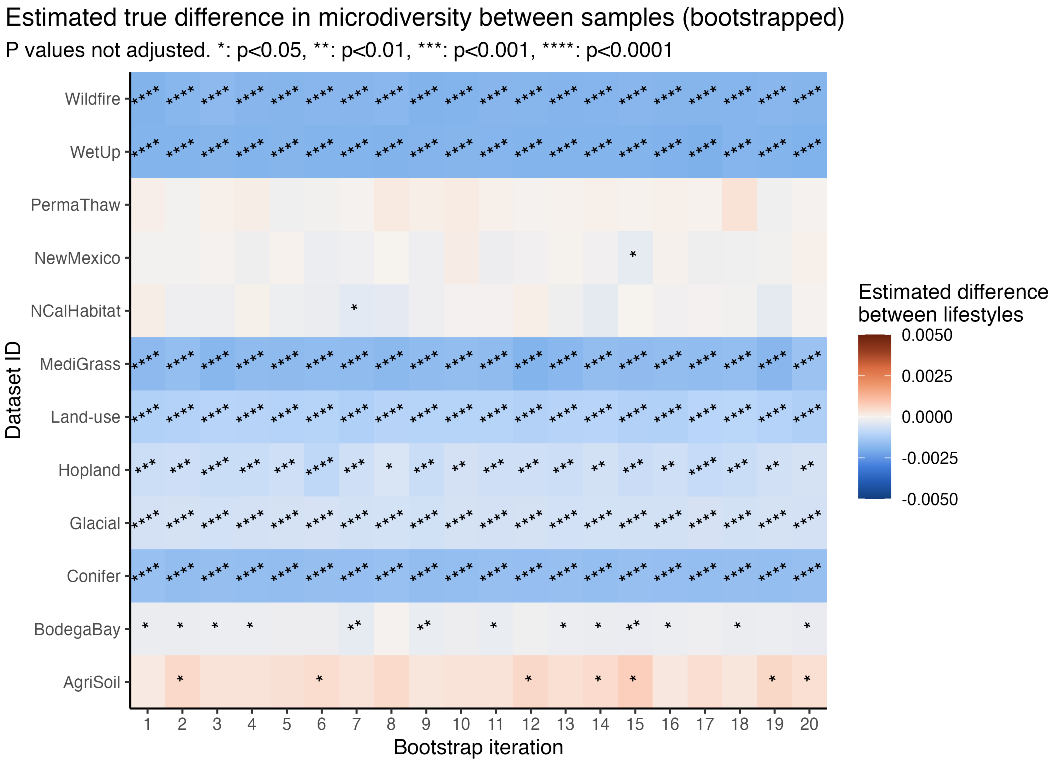


**Supplemental Figure 4. Bootstrapping shows robust lifestyle-microdiversity association**. Downsampled to lowest abundance lifestyle, Wilcoxon rank sum test for statistical association with asterisks indicating significance. Pattern across bootstrapping iterations is very similar to the non-downsampled results.

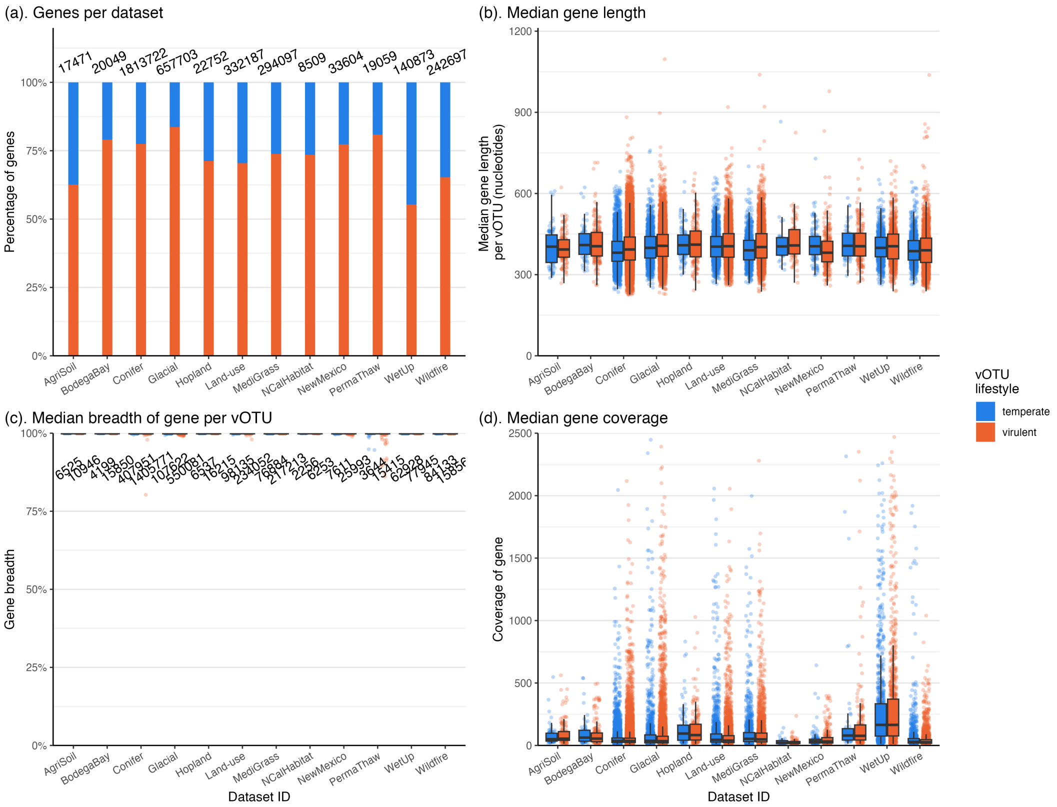


**Supplemental Figure 5. General statistics of genes**. All genes shown originate from vOTUs that passed the quality thresholds. (**A**). Number of genes per dataset and ratio of lifestyles of the vOTU they originate from. These ratios are similar to the ratios of the vOTUs. (**B**). Median length of genes, split by lifestyle. No patterns are visible due to vOTU lifestyle. (**C**). Median breadth of mapping of genes. These values are higher than those of vOTU breadth. (**D**). Median gene coverage (times). These values are similar to those of vOTU coverage.

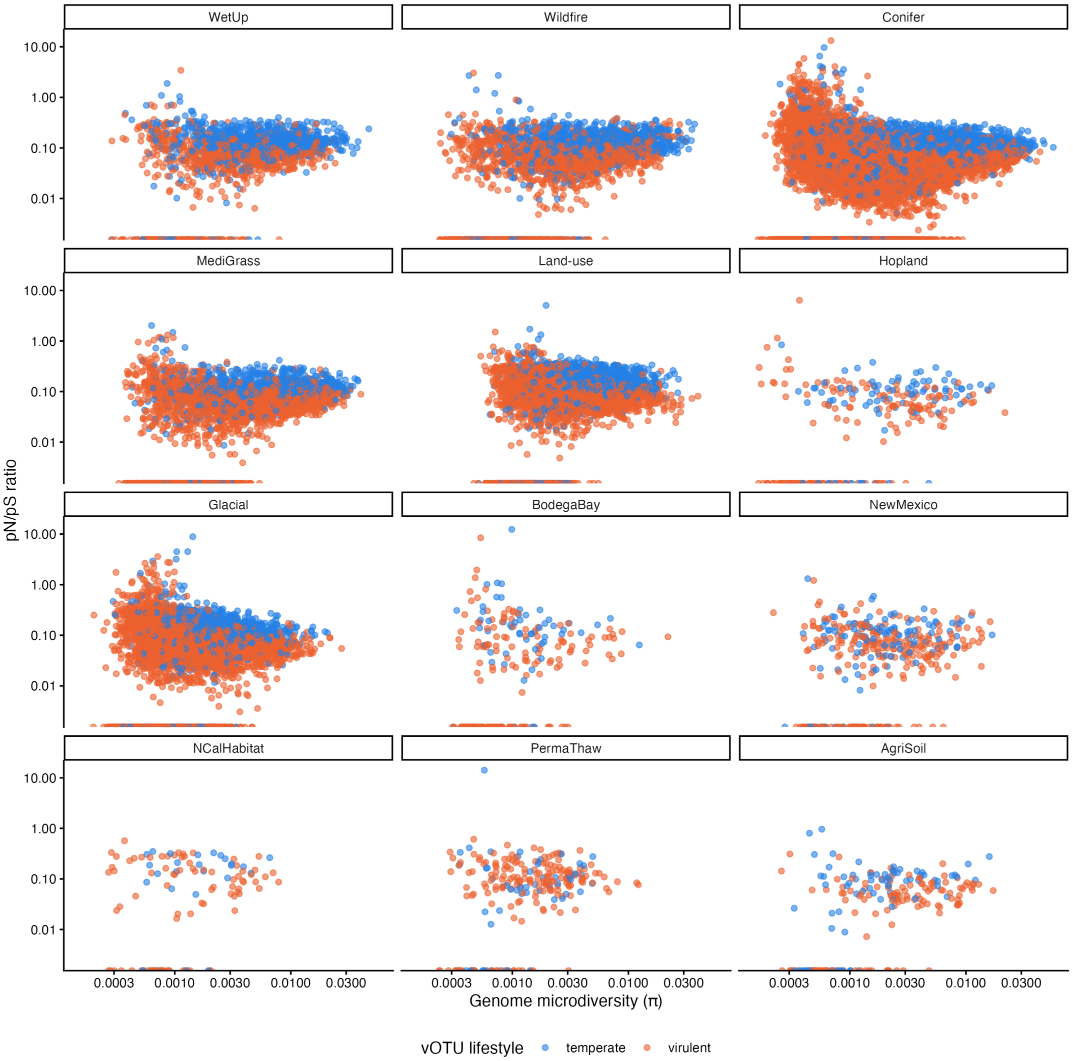


**Supplemental Figure 6. pN/pS versus microdiversity plotted per dataset**.

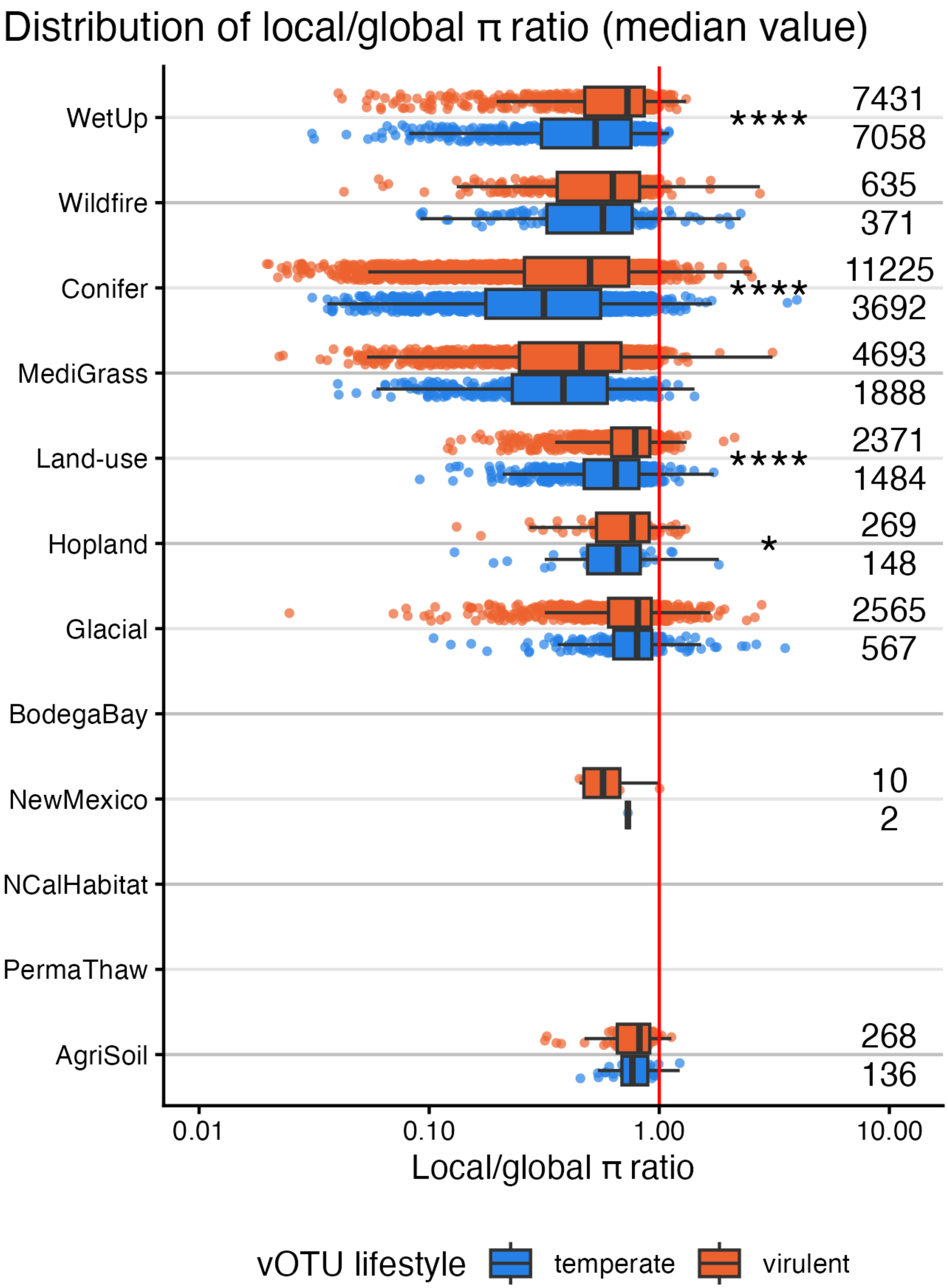


**Supplemental Figure 7. Ratio between local versus global diversity per dataset and lifestyle**. Temperate phages show generally lower ratio between local and global diversity, implying libraries differ more between each other than within compared to virulent phages.

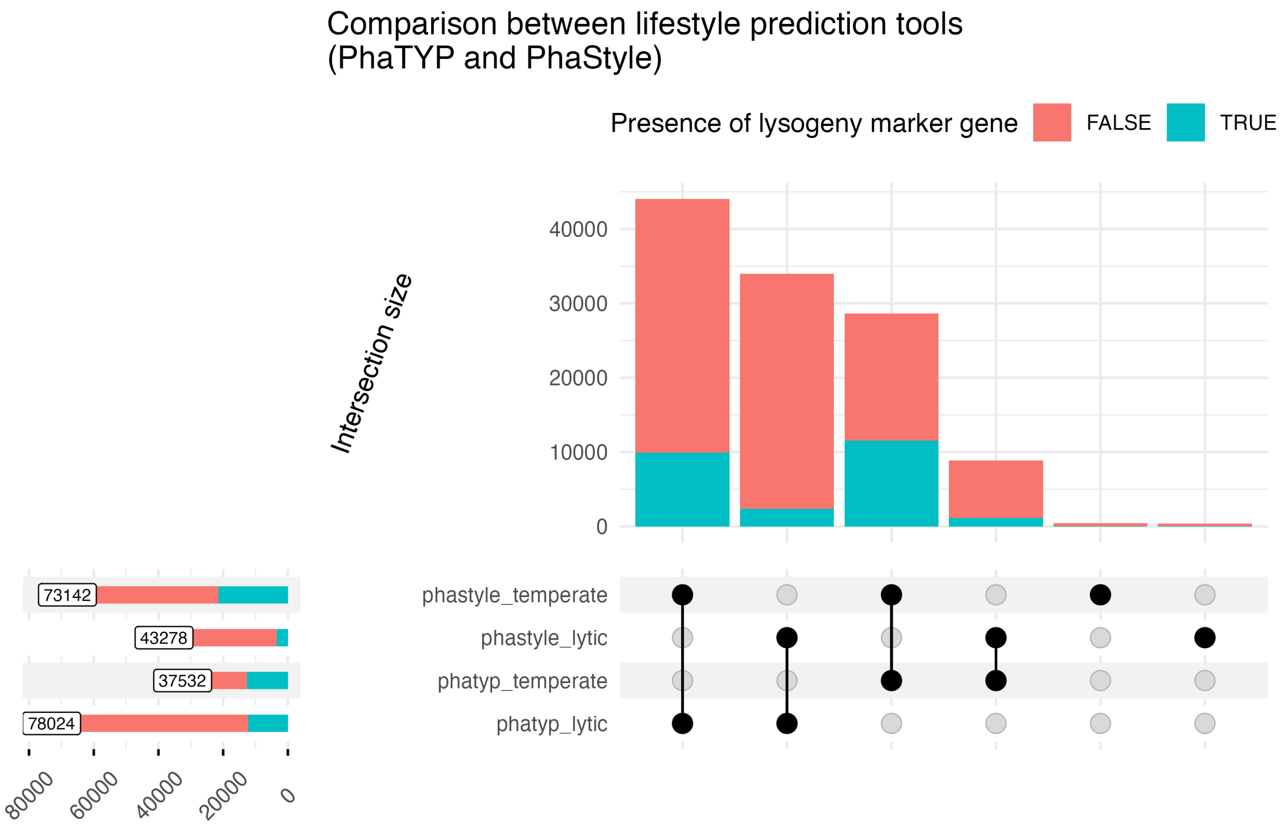


**Supplemental Figure 8. Upset plot showing lifestyle prediction groups per tool**.

#### Supplemental Tables

**Supplemental Table 1. Mapping efficiencies of all seventeen datasets**. The "total reads" column contains the combined number of reads from all included libraries, before mapping. The mapping efficiency, combined with the number of reads, supports the decision to only include viromes in the data analysis, as these tend to have a high mapping efficiency.

| Dataset ID | Libraries used for mapping | Mapping efficiency viromes (avg.) | Mapping efficiency metagenomes (avg.) | Total reads |
|---|---|---:|---:|---:|
| AgriSoil | 15 (v) 16 (m) | 27.73% | 0.42% | 518 499 214 |
| BodegaBay | 63 (v) 63 (m) | 11.30% | 2.71% | 7 507 979 642 |
| Conifer | 90 (v) | 64.81% | - | 3 371 778 675 |
| Glacial | 24 (v) | 55.79% | - | 1 698 161 279 |
| Hopland | 44 (v) | 3.24% | - | 2 671 824 516 |
| Intertidal | 96 (m) | - | 0.96% | 7 169 436 695 |
| Land-use | 50 (v) | 37.07% | - | 3 168 200 614 |
| MediGrass | 59 (v) | 30.59% | - | 2 777 968 364 |
| NCalHabitat | 30 (v) | 0.81% | - | 1 226 013 651 |
| NewMexico | 11 (v) | 6.23% | - | 983 697 171 |
| PermaThaw | 20 (v) | 20.66% | - | 2 158 529 983 |
| Rhizo | 12 (v) | 50.18% | - | 97 558 067 |
| Spruce | 10 (v) | 3.28% | - | 486 488 059 |
| UKagri | 15 (v) | 19.78% | - | 16 555 479 |
| Watershed | 47 (m) | - | 1.23% | 7 871 535 257 |
| WetUp | 152 (v) | 21.25% | - | 6 487 777 958 |
| Wildfire | 39 (v) | 48.82% | - | 1 452 099 335 |

**Supplemental Table 2. Statistical results regarding association between lifestyle and quality metrics used for filtering**. Reference point with regard to estimated true difference is temperate lifestyle (e.g. for length, virulent phages have a longer vOTU sequence). Tested using two-sided Wilcoxon rank sum test.

| Metric | Dataset ID | p-value | Estimated true difference |
|---|---|---|---|
| Length | AgriSoil | **5.89e-03** | 3 719 |
| Length | BodegaBay | **1.26e-04** | 7 012 |
| Length | Conifer | **1.43e-36** | 1 947 |
| Length | Glacial | **6.65e-16** | 3 113 |
| Length | Hopland | **1.07e-06** | 10 038 |
| Length | Land-use | **3.62e-50** | 4 516 |
| Length | MediGrass | **2.27e-32** | 4 420 |
| Length | NCalHabitat | **4.41e-02** | 5 490 |
| Length | NewMexico | **3.65e-10** | 11 995 |
| Length | PermaThaw | 7.00e-02 | - |
| Length | WetUp | **6.86e-24** | 5 101 |
| Length | Wildfire | **7.21e-25** | 5 771 |
| Completeness | AgriSoil | **1.98e-02** | 0.0000369 |
| Completeness | BodegaBay | 2.67e-01 | - |
| Completeness | Conifer | **1.55e-41** | 0.0000317 |
| Completeness | Glacial | **4.83e-21** | 0.0000938 |
| Completeness | Hopland | 3.10e-01 | - |
| Completeness | Land-use | **4.01e-18** | 0.0000284 |
| Completeness | MediGrass | **2.28e-04** | 0.0000576 |
| Completeness | NCalHabitat | **1.27e-04** | 2.50 |
| Completeness | NewMexico | 2.38e-01 | - |
| Completeness | PermaThaw | 3.70e-01 | - |
| Completeness | WetUp | **6.29e-03** | 0.0000477 |
| Completeness | Wildfire | **8.45e-16** | 0.0000659 |
| Coverage | AgriSoil | 7.99e-1 | - |
| Coverage | BodegaBay | 3.74e-1 | - |
| Coverage | Conifer | **2.79e-4** | 1.28 |
| Coverage | Glacial | **1.01e-2** | 1.96 |
| Coverage | Hopland | 9.76e-1 | - |
| Coverage | Land-use | **2.48e-9** | 5.76 |
| Coverage | MediGrass | **1.17e-3** | 4.60 |
| Coverage | NCalHabitat | 9.46e-1 | - |
| Coverage | NewMexico | 2.56e-1 | - |
| Coverage | PermaThaw | 9.62e-1 | - |
| Coverage | WetUp | 9.42e-1 | - |
| Coverage | Wildfire | **1.92e-6** | 3.06 |
| Breadth | AgriSoil | 8.90e-02 | - |
| Breadth | BodegaBay | 7.63e-01 | - |
| Breadth | Conifer | **1.54e-07** | 0.0000015 |
| Breadth | Glacial | **3.50e-02** | 0.0000023 |
| Breadth | Hopland | 4.66e-01 | - |
| Breadth | Land-use | **3.15e-14** | -0.0000106 |
| Breadth | MediGrass | 5.79e-02 | - |
| Breadth | NCalHabitat | 4.91e-01 | - |
| Breadth | NewMexico | 4.08e-01 | - |
| Breadth | PermaThaw | 8.60e-01 | - |
| Breadth | WetUp | 3.74e-01 | - |
| Breadth | Wildfire | **2.16e-03** | -0.0000017 |

**Supplemental Table 3. General genome statistics per dataset**. * The metrics "Length", "Coverage", "Mapping breadth", "Completeness" and "Nucleotide diversity" are all averages per dataset.

| Dataset ID | # of vOTUs | Length* | Coverage* | Mapping breadth* | Completeness* | Nucleotide diversity* |
|---|---|---|---|---|---|---|
| AgriSoil | 224 | 50 660 | 168 | 1.00 | 97.6 | 0.0029 |
| BodegaBay | 264 | 52 060 | 154 | 1.00 | 96.4 | 0.0014 |
| Conifer | 22 335 | 55 213 | 61 | 1.00 | 98.1 | 0.0037 |
| Glacial | 6 517 | 68 704 | 128 | 1.00 | 97.4 | 0.0018 |
| Hopland | 280 | 55 364 | 165 | 1.00 | 97.0 | 0.0026 |
| Land-use | 4 543 | 49 924 | 121 | 0.999 | 96.7 | 0.0040 |
| MediGrass | 3 557 | 55 730 | 113 | 1.00 | 98.1 | 0.0058 |
| NCalHabitat | 121 | 50 126 | 30 | 1.00 | 96.3 | 0.0017 |
| NewMexico | 399 | 57 154 | 63 | 0.999 | 94.8 | 0.0026 |
| PermaThaw | 245 | 55 955 | 412 | 0.987 | 95.3 | 0.0018 |
| WetUp | 1 764 | 53 212 | 454 | 1.00 | 97.6 | 0.0054 |
| Wildfire | 2 922 | 55 066 | 62 | 1.00 | 97.0 | 0.0045 |

**Supplemental Table 4. vOTU length and nucleotide diversity statistics per dataset per lifestyle**. Clear differences in the number of vOTUs that pass quality thresholds are visible between studies. * The metrics "Length" and "Nucleotide diversity" are averages per dataset.

| Dataset ID | Lifestyle | # of vOTUs | Length* | Min. length | Max. length | Nucleotide diversity* |
|---|---|---|---|---|---|---|
| AgriSoil | temperate | 85 | 48 485 | 21 170 | 259 025 | 0.0023 |
| AgriSoil | virulent | 139 | 51 990 | 13 770 | 153 522 | 0.0032 |
| BodegaBay | temperate | 65 | 42 813 | 14 015 | 100 963 | 0.0015 |
| BodegaBay | virulent | 199 | 55 080 | 13 723 | 195 821 | 0.0014 |
| Conifer | temperate | 5 281 | 50 144 | 14 362 | 272 124 | 0.0059 |
| Conifer | virulent | 17 054 | 56 782 | 12 247 | 462 798 | 0.0030 |
| Glacial | temperate | 1 381 | 51 397 | 13 289 | 198 996 | 0.0024 |
| Glacial | virulent | 5 136 | 73 358 | 12 835 | 676 544 | 0.0016 |
| Hopland | temperate | 94 | 45 115 | 12 996 | 89 616 | 0.0033 |
| Hopland | virulent | 186 | 60 543 | 12 966 | 316 021 | 0.0022 |
| Land-use | temperate | 1 440 | 44 774 | 13 024 | 164 636 | 0.0048 |
| Land-use | virulent | 3 103 | 52 314 | 10 069 | 373 888 | 0.0036 |
| MediGrass | temperate | 1 003 | 49 228 | 13 233 | 150 206 | 0.0077 |
| MediGrass | virulent | 2 554 | 58 284 | 12 793 | 438 185 | 0.0050 |
| NCalHabitat | temperate | 32 | 46 905 | 20 121 | 95 220 | 0.0017 |
| NCalHabitat | virulent | 89 | 51 284 | 15 235 | 103 915 | 0.0017 |
| NewMexico | temperate | 119 | 42 863 | 13 699 | 88 220 | 0.0026 |
| NewMexico | virulent | 280 | 63 228 | 13 967 | 362 520 | 0.0027 |
| PermaThaw | temperate | 53 | 48 014 | 31 941 | 92 431 | 0.0017 |
| PermaThaw | virulent | 192 | 58 148 | 17 908 | 288 904 | 0.0018 |
| WetUp | temperate | 862 | 47 423 | 17 026 | 144 690 | 0.0069 |
| WetUp | virulent | 902 | 58 745 | 13 525 | 335 613 | 0.0039 |
| Wildfire | temperate | 1 109 | 48 664 | 12 788 | 147 325 | 0.0063 |
| Wildfire | virulent | 1 813 | 58 982 | 11 813 | 357 464 | 0.0034 |

**Supplemental Table 5. Statistical results regarding association between lifestyle and microdiversity, tested using multiple lifestyle prediction methods**. Reference point is temperate lifestyle. Default settings include using PhaTYP, PhaStyle and DIAMOND BLASTp. P-values below 0.05 have been marked in bold. 95% confidence intervals that do not span zero and belong to a p-value below 0.05 are also marked in bold.

| Lifestyle tools | Dataset ID | f-statistic | p-value | 95% CI low | 95% CI high |
|---|---|---:|---:|---|---|
| Default | AgriSoil | 4 902 | **3.27e-002** | **-0.001021** | **-0.000052** |
| Default | BodegaBay | 7 878 | **8.34e-003** | **0.000036** | **0.000367** |
| Default | Conifer | 61 776 988 | **0** | **0.001532** | **0.001736** |
| Default | Glacial | 4 939 695 | **1.36e-111** | **0.000539** | **0.000648** |
| Default | Hopland | 11 287 | **6.99e-005** | **0.000307** | **0.001173** |
| Default | Land-use | 2 917 652 | **5.32e-062** | **0.000963** | **0.001206** |
| Default | MediGrass | 1 633 009 | **2.16e-037** | **0.001441** | **0.002050** |
| Default | NCalHabitat | 1 504 | 6.40e-001 | -0.000249 | 0.000431 |
| Default | NewMexico | 17 462 | 4.47e-001 | -0.000181 | 0.000372 |
| Default | PermaThaw | 4 874 | 6.40e-001 | -0.000348 | 0.000230 |
| Default | WetUp | 522 570 | **6.38e-036** | **0.001589** | **0.002279** |
| Default | Wildfire | 1 386 013 | **2.53e-066** | **0.001633** | **0.002058** |
| PhaStyle only | AgriSoil | 42 431 | **1.00e- 2** | **-0.000605** | **-0.0000722** |
| PhaStyle only | BodegaBay | 100 810 | **6.56e- 3** | **0.0000112** | **0.000152** |
| PhaStyle only | Conifer | 477 079 424 | **2.03e-273** | **0.000398** | **0.000488** |
| PhaStyle only | Glacial | 34 044 640 | **3.04e-116** | **0.000247** | **0.000279** |
| PhaStyle only | Hopland | 86 037 | **6.59e- 3** | **0.0000472** | **0.000337** |
| PhaStyle only | Land-use | 25 219 741 | **4.04e-108** | **0.000693** | **0.000831** |
| PhaStyle only | MediGrass | 14 687 043 | **8.10e- 51** | **0.000800** | **0.00112** |
| PhaStyle only | NCalHabitat | 15 649 | 6.30e- 1 | -0.000121 | 0.000210 |
| PhaStyle only | NewMexico | 156 359 | 1.28e- 1 | -0.0000413 | 0.000259 |
| PhaStyle only | PermaThaw | 54 548 | 7.59e- 1 | -0.000139 | 0.000161 |
| PhaStyle only | WetUp | 2 985 555 | **1.92e- 37** | **0.000903** | **0.00129** |
| PhaStyle only | Wildfire | 8 935 581 | **4.64e- 69** | **0.000863** | **0.00110** |
| PhaTYP only | AgriSoil | 62 265 | 5.19e- 1 | -0.000114 | 0.000200 |
| PhaTYP only | BodegaBay | 95 758 | **4.12e- 2** | **-0.0000119** | **0.000130** |
| PhaTYP only | Conifer | 388 073 918 | **4.11e-33** | **0.000114** | **0.000169** |
| PhaTYP only | Glacial | 26 272 742 | **3.01e-29** | **0.000101** | **0.000144** |
| PhaTYP only | Hopland | 91 665 | **4.44e- 3** | **0.0000670** | **0.000436** |
| PhaTYP only | Land-use | 23 607 107 | **1.55e-22** | **0.000286** | **0.000409** |
| PhaTYP only | MediGrass | 12 980 938 | **4.14e-10** | **0.000292** | **0.000572** |
| PhaTYP only | NCalHabitat | 14 757 | 9.58e- 1 | -0.000200 | 0.000164 |
| PhaTYP only | NewMexico | 143 217 | 6.63e- 1 | -0.000126 | 0.000177 |
| PhaTYP only | PermaThaw | 53 278 | **3.22e- 2** | **-0.00000592** | **0.000331** |
| PhaTYP only | WetUp | 3 220 506 | **8.51e- 7** | **0.000224** | **0.000566** |
| PhaTYP only | Wildfire | 9 002 495 | **2.21e-26** | **0.000539** | **0.000766** |
| PhaStyle + PhaTYP | AgriSoil | 14 866 | 5.27e- 2 | -0.000699 | -0.0000168 |
| PhaStyle + PhaTYP | BodegaBay | 27 154 | **1.70e- 3** | **0.0000309** | **0.000226** |
| PhaStyle + PhaTYP | Conifer | 152 551 464 | **2.89e-210** | **0.000533** | **0.000622** |
| PhaStyle + PhaTYP | Glacial | 11 358 986 | **4.19e-102** | **0.000289** | **0.000357** |
| PhaStyle + PhaTYP | Hopland | 27 838 | **3.21e- 4** | **0.000153** | **0.000673** |
| PhaStyle + PhaTYP | Land-use | 8 002 181 | **2.13e- 93** | **0.000875** | **0.00107** |
| PhaStyle + PhaTYP | MediGrass | 4 326 644 | **3.64e- 43** | **0.00107** | **0.00152** |
| PhaStyle + PhaTYP | NCalHabitat | 4 493 | 7.36e- 1 | -0.000187 | 0.000264 |
| PhaStyle + PhaTYP | NewMexico | 44 012 | 2.05e- 1 | -0.0000979 | 0.000347 |
| PhaStyle + PhaTYP | PermaThaw | 17 735 | 1.20e- 1 | -0.0000425 | 0.000371 |
| PhaStyle + PhaTYP | WetUp | 1 020 429 | **2.84e- 30** | **0.00106** | **0.00154** |
| PhaStyle + PhaTYP | Wildfire | 3 077 060 | **6.53e- 71** | **0.00119** | **0.00157** |

**Supplemental Table 6. Statistical results regarding difference between pN/pS of genes** between temperate and virulent phages, as tested using a two-sided Wilcoxon rank sum test.

| Dataset ID | f-statistic | p-value | 95% CI low | 95% CI high |
|---|---|---|---|---|
| AgriSoil | 4 180 645 | **1.40e-25** | 0.0154 | 0.0333 |
| BodegaBay | 1 041 598 | **8.40e-13** | 0.0079 | 0.0397 |
| Conifer | 57 331 369 800 | **0** | 0.0279 | 0.0290 |
| Glacial | 2 672 129 586 | **0** | 0.0313 | 0.0351 |
| Hopland | 6 489 458 | **6.60e-50** | 0.0288 | 0.0405 |
| Land-use | 2 593 954 210 | **0** | 0.0431 | 0.0467 |
| MediGrass | 2 571 859 234 | **0** | 0.0397 | 0.0423 |
| NCalHabitat | 656 806 | **2.10e- 5** | 0.0000125 | 0.0334 |
| NewMexico | 8 798 160 | **3.60e-27** | 0.0110 | 0.0276 |
| PermaThaw | 2 063 826 | 2.47e- 1 | -0.0000664 | 0.0000678 |
| WetUp | 600 436 849 | **0** | 0.0454 | 0.0494 |
| Wildfire | 1 499 329 808 | **0** | 0.0498 | 0.0531 |
